# Transcriptomic analysis of mdx mouse muscles reveals a signature of early human Duchenne muscular dystrophy

**DOI:** 10.1101/2021.07.16.452553

**Authors:** Evelyn Ralston, Gustavo Gutierrez-Cruz, Aster Kenea, Stephen R. Brooks

**Affiliations:** Light Imaging Section, Office of Science and Technology, National Institute of Arthritis and Musculoskeletal and Skin Diseases (NIAMS), NIH, Bethesda, MD 20892, USA; Genomic Technology Section, Office of Science and Technology, National Institute of Arthritis and Musculoskeletal and Skin Diseases (NIAMS), NIH, Bethesda, MD 20892, USA; Biodata Mining and Discovery Section, Office of Science and Technology, National Institute of Arthritis and Musculoskeletal and Skin Diseases (NIAMS), NIH, Bethesda, MD 20892, USA

**Keywords:** muscle, Duchenne, *mdx*, mouse model, RNA-seq

## Abstract

The *mdx* mouse (C57BL/10ScSn-*DMD^mdx^*/J) is the oldest model of Duchenne muscular dystrophy (DMD). *Mdx* remains popular and has not been replaced by newer mouse models, despite criticisms that *mdx* has a nearly normal lifespan and mild pathology while DMD remains a severe, fatal disease. At some point we noticed that the absence of *mdx* RNA-seq data limited our ability to assess the results of physiological work on the mouse model and to compare these results to human genetic data [1]. We carried out RNA-seq analysis of wild-type and *mdx* mice of 2 and 5 months of age, using three hindlimb muscles per mouse: the *flexor digitorum brevis (FDB)*, the *extensor digitorum longus (EDL)* and the *soleus (SOL)*, with a total of 55 samples. We then mined the data and found that each of the three muscles is a valid experimental model for DMD-related mouse work, even the FDB, despite a delayed pathology development. We also show that the *mdx* mouse muscles are enriched in metabolic, developmental, regenerational and structural pathways that have been found to be the “disease signature” of DMD in young and presymptomatic subjects [38, 39]. Additionally, we show that healthy human muscle fiber microtubules present the grid-like organization found in control rodents but perturbed in the *mdx* mouse. We conclude that the *mdx* mouse appropriately mimics the early stages of DMD, with its microtubule defects signaling fiber regeneration [35]. We hope that these results may contribute to a better understanding of the failure of regeneration as DMD progresses.

## 1. INTRODUCTION^1^

Dystrophin is the protein missing in boys affected by the X-linked Duchenne muscular dystrophy (DMD). Both dystrophin [2] and its gene, *DMD* [3], were characterized 34 years ago. Regrettably, there is still no cure for the disease. DMD remains fatal even if the lifespan of those affected has been prolonged by anti-inflammatory drugs and patient care measures. A treatment based on exon-skipping to produce a shortened but functional dystrophin, was approved in 2016 [4–5] but initial assessment of the results was only mildly encouraging [6]. Furthermore, exon-skipping reagents can only help DMD-affected boys with the appropriate mutation [7–8]. Several other approaches appear promising [9] but basic research remains essential and animal models play a crucial role. The golden retriever dog model (GRMD), discovered in 1980 before the *mdx* mouse and the *DMD* gene, is increasingly appreciated as a pre-clinical model [10] but mice remain the first tool for early large-scale studies.

The C57BL/10 *mdx* mouse (C57BL/10ScSn-*DMD^mdx^*/J) was the first mouse model of DMD. A spontaneous mutation prevents dystrophin transcription and translation [11–12]. It is a true genetic model of the human disease. Pathology starts at 4 weeks of age with a burst of muscle fiber degeneration and regeneration that later slow down. Most muscles show compensatory hypertrophy from 10 to 40 weeks of age [13] and some atrophy thereafter with inflammation and cycles of fiber death and regeneration. *Mdx* mouse muscles undergo membrane damage upon muscle stretching, excessive production of reactive oxygen species (ROS) linked to an increase in the NADPH oxidase NOX2 and fibrosis, all hallmarks of DMD [1,14]. Furthermore, *mdx* muscles show structural abnormalities such as an increase in split fibers [13] and a disordered microtubule pattern [15–16]. Quantitation of such abnormalities has helped evaluate *mdx* mouse lines either rescued or made sicker by transgenic expression or ablation of other proteins. Microtubule pattern analysis has been used for investigating the contribution of various motifs of the very large (427 kDa) dystrophin protein [17–19]. Studies of the *mdx*-Fiona mouse, which overexpresses the dystrophin homolog utrophin, has led to clinical trials of utrophin overexpression [20–21] and small molecules that modulate utrophin are being successfully developed [22].

Despite these achievements, the *mdx* mouse has been criticized: its lifespan is close to that of the WT mouse and its pathology is mild compared to that of the human disease [23] with inflammation as its dominant pathology [24]. Several other mouse lines have been developed [25–29]. These lines are affected faster and more severely and may have a shorter lifespan than *mdx* mice. Nevertheless, the *mdx* mouse remains widely used [30].

Our interest in the *mdx* mouse transcriptome was stimulated by a publication that causally linked microtubule disorganization of the *mdx* mouse and human DMD pathology [1]. We and others had found that *mdx* muscle microtubules are disorganized compared to WT ones [15–16] but the implications of this disorganization were not clear. The cited study [1] suggested that *mdx* microtubules were toxic: depolymerization of microtubules prevented ROS production and Ca^2+^ uptake by *mdx* muscle fibers stretched *ex vivo*. When WT fibers were stretched, there was no –or much less– ROS produced. In support for a microtubule role, RNA sequencing (RNA-seq) of DMD vs. normal (NORM) human muscle biopsies highlighted transcriptional changes in genes associated with ROS production and microtubule composition [1]. However, some data was lacking to further evaluate these striking results.

First, there was no RNA-seq of *mdx* mice for a comparison against human muscles and mouse microarray studies only covered a limited number of microtubule components. Secondly, we had no information on the organization of human compared to mouse muscle microtubules. Thirdly, the cited work was carried out on the *flexor digitorum brevis* (FDB) muscle, a small muscle of the mouse foot plant. The FDB is convenient to use because it is easy to digest with collagenase [31], producing single fibers that can be plated and investigated fixed or live [32–35]. But DMD affects core upper muscles such as the diaphragm earlier and more severely than lower limb muscles and the FDB is as far from the upper core muscles as possible in a mouse. There were thus reasons to ask whether the FDB is as good a model of DMD as other muscles such as the *extensor digitorum longus* (EDL) muscle, frequently used for physiological studies [36].

We therefore carried out RNA-seq on *mdx* and WT mouse muscles. We choose three muscles: the EDL, the FDB, and the *soleus* (SOL). Muscle fiber type composition contributes importantly to muscle physiology [37] and these three muscles are representative: the EDL is especially rich in fast fibers, the FDB in fast intermediate fibers and the SOL in slow fibers. We selected the ages of 2 and 5 months, because the fiber stretching experiments [1] had shown a strong difference between 5 month old and younger animals. A total of 55 samples were sequenced and analyzed by comparison between genotypes and ages and further compared with DMD data, starting with microarray analyses of young (5-7 years) [38] and presymptomatic (younger than 2 years) [39] DMD cases and continuing with our own analysis of the published DMD data from [1]. We show that the *mdx* mouse best models early stages of DMD; we validate the use of the FDB muscle; and we find that human muscle fiber microtubules do have a grid-like organization as has been found in rodent muscles.

## 2. MATERIALS & METHODS

### 2.1. Animals

C57BL/10ScSn/J and C57BL/10ScSn-Dmd/J mice (hereafter referred to as wt and *mdx*, respectively) were obtained from the Jackson Laboratory (Bar Harbor, ME) in groups of 3 to 6 at the age of 1-2 mo and housed in the NIAMS Bldg. 50 Animal Facility until they were 2 or 5 months-old. All animals were male since the dystrophin-null mutation is X-linked and only affects males, in humans as in mice.

### 2.2. Muscle collection and total RNA extraction

When mice had reached the age of 2 or 5 months, they were killed with CO_2_, one at a time. *EDL, FDB, and SOL* muscles were collected and immediately dropped into *RNAlater* (Ambion, ThermoFisher Scientific, Carlsbad, CA). EDL and SOL were then immediately chopped into small fragments while FDB was first cleaned under a dissection microscope to remove tendons and large blood vessels and then cut into fragments. All instruments and surfaces in contact with the muscles had been pretreated with RNase away (Molecular BioProducts, Inc., San Diego, CA). From then on, all samples were handled in parallel. Muscle fragments were transferred to 1 ml Trizol reagent (InVitrogen, ThermoFisher Scientific) in 2ml tubes containing 1.4 mm ceramic beads (VWR, Rador, PA) for pulverization in a Precellys homogenizer (Bertin, France) with 2 rounds at 6,000 rpm for 15 secs each. After addition of chloroform-methanol, the muscle extracts were spun for 15 min in a cooled Eppendorf table-top centrifuge, the aqueous phase was recovered, and RNA was extracted with a PureLink RNA Mini Kit and treated with PureLink DNase (InVitrogen ThermoFisher Scientific). RNA was quantitated and its quality verified in an Agilent 2100 bioanalyzer (Agilent Technologies, Waldbronn, Germany) and kept at −80°C for further RNA-seq library preparation. Only samples with a RIN >8.5 were retained.

### 2.3. RNA-Seq library preparation

From the total RNA, we then isolated, fragmented and primed the mRNA with NEB Next Poly (A) mRNA Magnetic Isolation Module (NEB #E7490, New England Biolabs, Ipswich, MA). The produced cDNA was quantitated with PicoGreen on a Perkin-Elmer Victor X3 fluorescence reader and kept at −80°C. cDNA library preparation for RNA-Seq was performed using the Mondrian SP kit (NuGEN Technologies, San Carlos CA). A total of 60 cDNA samples underwent RNA-seq. RNA sequencing was performed in the NIAMS Genome Core Facility at the NIH.

### 2.4. RNA-Seq analysis

RNA-Seq libraries were run on an Illumina HiSeq2000 sequencer (Illumina, San Diego, CA) as 50 base, single end runs. Samples were demultiplexed and converted to FastQ files using Casava version 1.8.2. FastQ files were mapped to mm10 using Tophat 2.1.0 and RPKM values were calculated using Partek Genomic Suites 6.6 (Partek, St Louis, MO). Principal Component Analysis (PCA), hierarchical clustering and heat map generation were performed using Partek GS. PCA analysis was performed on RPKM data with 12,000 genes (filtered for expressed genes, i.e. max RPKM > 1). Five samples did not pass our quality control and were excluded from further analysis, leaving 55 samples as follows.

| *muscle* | *genotype* | *age (months)* | *number of samples* |
| --- | --- | --- | --- |
| EDL | WT | 2 | 4 |
|  |  | 5 | 4 |
|  | mdx | 2 | 4 |
|  |  | 5 | 6 |
| SOL | WT | 2 | 4 |
|  |  | 5 | 4 |
|  | mdx | 2 | 4 |
|  |  | 5 | 7 |
| FDB | WT | 2 | 4 |
|  |  | 5 | 4 |
|  | mdx | 2 | 3 |
|  |  | 5 | 7 |

Hierarchical clustering was performed on 2503 genes (filtered using Max RPKM > 1 and CV > 0.3). ANOVA comparisons were performed on log_2_ transformed RPKM values (with a 0.1 offset) and false discovery rate (q-value) calculated using Partek GS.

### 2.5. Human RNA-Seq analysis

Human RNA-Seq results were downloaded from Table S2 in [1] (https://stke.sciencemag.org/content/suppl/2012/08/03/5.236.ra56.DC1). Six DMD samples had been obtained from subjects diagnosed with DMD who were 11 mo, and 4, 5 (two samples), 7, and 8 years old. The values in these tables are log_2_ transformed expression values. One of the control samples, NORM3223, was eliminated as an outlier. ANOVA comparison was performed as above using Partek GS.

### 2.6. Pathway Enrichment Analysis

Gene lists were filtered keeping only genes for which at least one sample group had an RPKM value ≥ 1, a fold change > 2 or < −2 and a false discovery rate (q-value) < 0.3. Combined pathway analysis was performed on 6 gene lists from *mdx* vs WT ANOVA in EDL, FDB and SOL muscles at 2 and 5 months and one gene list from DMD results [1]. We included mouse genes that were 2-fold up and with a false discovery rate (q-value) < 0.3. DMD results were filtered at 5-fold up. These 7 gene lists were submitted to Metascape.org (https://pubmed.ncbi.nlm.nih.gov/30944313/) and integrated pathway analysis reported.

### 2.7. Plots

Venn diagrams (Fig. 3) were drawn with Venny 2.1.0 (Oliveros, 2007-15; https://bioinfogp.cnb.csic.es/tools/venny/). Bubble plots (Fig. 5) and the directionality plot (Fig. 7) were done in JMP 14 (SAS, Cary, NC).

### 2.8. Human muscle fiber immunofluorescence

Bundles of human muscle fibers containing up to 20 fibers each were provided by Drs. Thorkil Ploug and the FINE group (Univ. of Copenhagen, Denmark), under an agreement of de-identification. The muscle biopsies were obtained from the middle portion of the vastus lateralis muscle from sedentary resting human subjects using the percutaneous needle technique with suction [40]. The part of the biopsy destined to be used for single fiber analysis was placed in Krebs–Ringer buffer containing 0.1% procaine hydrochloride for 2–3 min and pinned down at resting length in a Petri dish coated with Sylgard 184 (Dow Corning, Midland, MI). It was then fixed with 2% depolymerized paraformaldehyde and 0.15% picric acid in 0.1 M phosphate buffer at room temperature for half an hour, followed by an additional 4.5 h in the fixative at 4°C, transfer to 50% glycerol in PBS and storage at −20°C [41]. Before immunofluorescence, fiber bundles were warmed up to room temperature and transferred from 50% glycerol to PBS. Single fibers were teased manually with fine forceps and immunofluorescence staining was carried out in multi-well dishes as previously reported for rat fibers [41]. Fibers were stained with the rat monoclonal YOL1/34 anti-a-tubulin (#100-1639) from Novus Biologicals (Littleton, CO) diluted 1:250 to label microtubules and with the mouse monoclonal F27 antibody against human GLUT4, a gift from Dr. T. Ploug. F27 was raised against a peptide corresponding to the 13 C-terminal amino-acids of human GLUT4 [40] and used diluted 1:50. Some samples were labeled only with the mouse monoclonal DM 1A anti-a-tubulin (#T9026) from Sigma-Aldrich (St-Louis, MO), diluted 1:500. Secondary antibodies were highly cross-adsorbed Alexa 488-conjugated goat anti-mouse IgG (InVitrogen, ThermoFisher, Waltham, MA) and Alexa 594-conjugated goat anti-rat IgG (Jackson ImmunoResearch, West Grove, PA). They were diluted 1:500. Control fibers were stained with secondary antibodies only. Fibers were finally counterstained with Hoechst 33342 (ThermoFisher) and mounted in Vectashield (Vector, Burlingame, CA). Images were collected on a Leica SP5 NLO confocal microscope (Leica Microsystems, Buffalo Grove, IL) with a 63x N.A. 1.4 objective. Images were cropped, linearly modified and assembled in Photoshop CC 2020 (Adobe, San Jose, CA). The original blue, green and red colors were modified using Photoshop image --> adjustments --> hue until proofing confirmed that the new colors could be distinguished in two cases of color blindness (Protanopia and Deuteranopia).

### 2.9. Microtubule directionality analysis

The Matlab-based TeDT software was developed in the NIAMS Light Imaging Section by Dr. Wenhua Liu [43] and is freely available for download from the Github database at https://github.com/ralstone/TeDT_microtubule_directionality. Data calculated in TeDT can easily be imported in MS Excel or in other software for plotting such as jmp14 as done here.

## 3. RESULTS

### 3.1. PCA and heatmap analyses show complex effects of muscle type, genotype and age on gene expression in mdx muscle

RNA-seq analysis of mouse muscles was carried out on a total of 60 samples and produced 35,680 transcripts for 24,453 genes. Five samples were identified as outliers and excluded from further analysis leaving 4 samples for each control group and from 3 to 7 samples for each *mdx* group (see methods). All results described here relate to gene-related differences without distinction between transcripts.

PCA analysis (Fig. 1) shows complete segregation of the three muscles (EDL in blue, FDB in green and SOL in yellow and red) due to differences in each of the three principal components. For each muscle, WT samples (lighter color hue) segregate neatly from *mdx* samples. Age (spheres vs. tetrahedrons) separates WT samples into two distinct groups (2- and 5-month-olds), but the separation is less clear for *mdx* samples.

**Figure 1:**
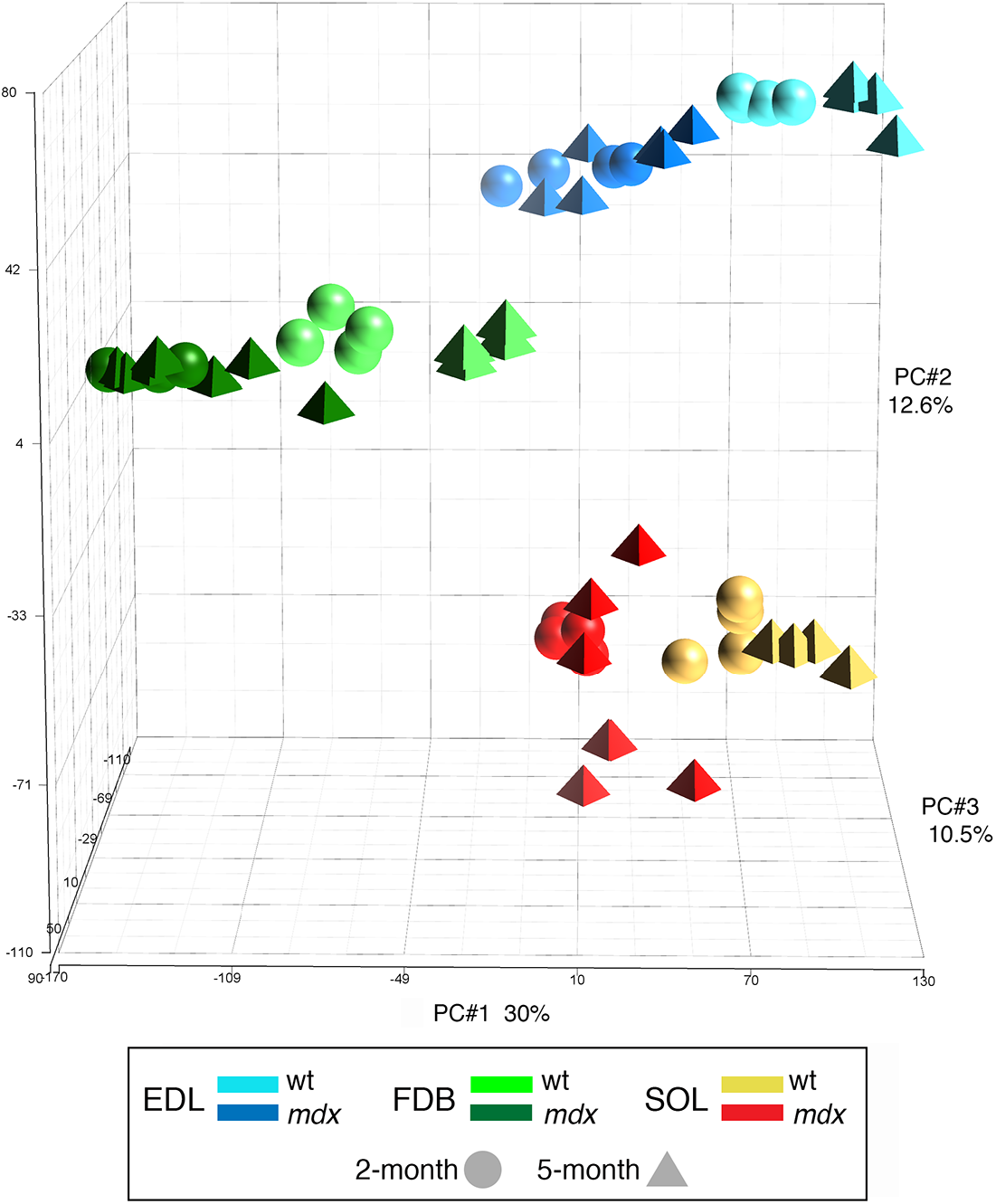
Muscle is the strongest discriminant in principal component analysis (PCA) of gene level RPKM data. All 55 samples and all expressed variable genes with at least one sample with RPKM>1 were subjected to PCA and plotted based on the first 3 principal components. FDB muscle is distinguished mostly by PC1, EDL and SOL muscles by PC2 and PC3. For each muscle, genotype (*mdx* vs WT) further segregates the samples and, finally, age segregates the samples within genotype, but less distinctly in 5- than in 2-month old groups.

Unguided hierarchical clustering (Fig. 2) shows the different ways in which genotype and age combine to affect gene expression. Some genes are predominantly expressed in one of the muscles (indicated above the plot as FDB≫, EDL≫, or SOL≫) with little difference between genotypes or ages. Other genes show increased expression in mdx compared to WT samples regardless of muscle ((indicated as mdx>WT), with some effect of age. Finally some of the genes with higher expression in the FDB muscle are affected by both genotype and age.

**Figure 2:**
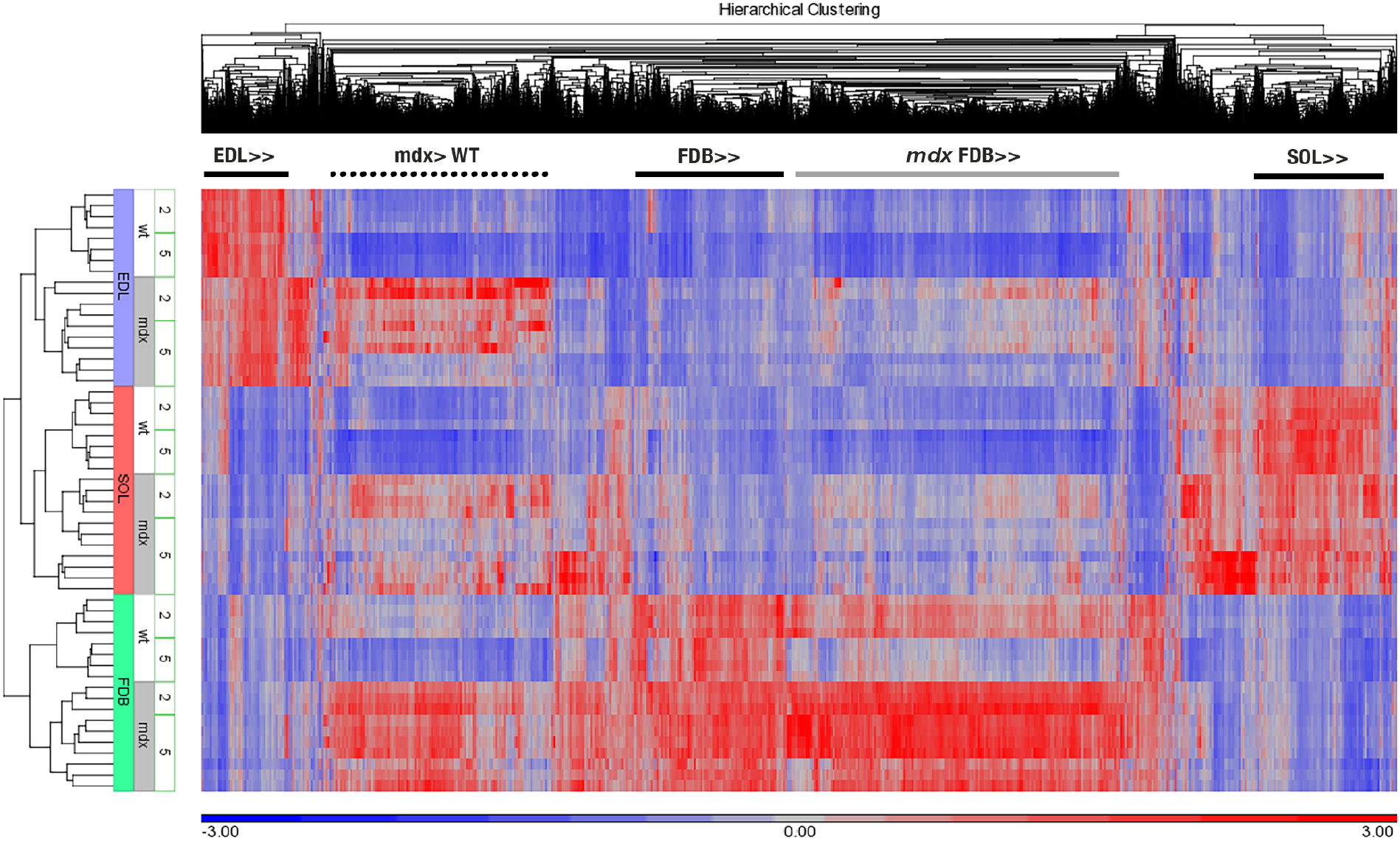
Hierarchical clustering of RPKM data highlights complex effects of muscle type, genotype and age on gene expression. Expressed variable genes were filtered to retain only those with at least one sample with RPKM>1 and a coefficient of variance across all 55 samples >0.3. All 55 samples and all retained genes (2,053) were then subjected to hierarchical clustering and the standardized RPKM values were plotted. Each standardized RPKM value is a standard deviation away from the mean for each gene across samples. Highest level clustering is by muscle, followed by genotype (*mdx* vs WT), and age. High expression shows up in red, low expression in blue. Some clusters of genes are predominantly expressed in one of the muscles (label EDL≫, FDB≫, or SOL≫ above the colored cells), regardless of genotype. One cluster of genes, in contrast, shows higher expression in *mdx* than in WT, regardless of the muscle. Others have a more complex expression pattern, for example the cluster of genes more highly expressed in the FDB but especially in the mdx genotype. Blocks of genes whose expression is muscle-dependent but genotype-independent are marked by a black line above the plot. A block of genes whose expression is, on the contrary, genotype-dependent but muscle-independent is indicated by a dotted bar. Additionally a large block of genes that show a different behavior and are found only in the FDB is indicated by a gray bar. Their expression is strongly increased in the *mdx* genotype but decreases during normal development (WT2>WT5).

We carried out ANOVA analysis of our mouse muscle RNA-seq results and of the human muscle RNA-seq results [1] available in the publication’s supplementary material (as described in the Methods). We filtered the results by fold change, retaining mouse genes whose expression is at least 2-fold higher in *mdx* than WT samples and human genes whose expression is at least 5-fold higher in DMD compared to control (NORM) samples. The results are plotted as Venn diagrams (Fig. 3) and listed in our Table S1. From 2 to 5 months the number of genes retained by our filter increases considerably more for the FDB (4.0 fold) than for the EDL (1.6 fold) or SOL (1.8 fold). Among the genes that are up in *mdx* samples, 180 are common to the three mouse muscles at 2 months (Fig. 3A) compared to 544 at 5 months. When mouse and human samples are compared (Fig. 3B), only 94 of the “up” genes are common to all at 2 months vs. 255 at 5 months. Thus, the dystrophic pathology is slower to manifest in the FDB than in the EDL and SOL muscles and the genes that are delayed in their response include some that are common to all three muscles as well as some specific to the FDB.

**Figure 3:**
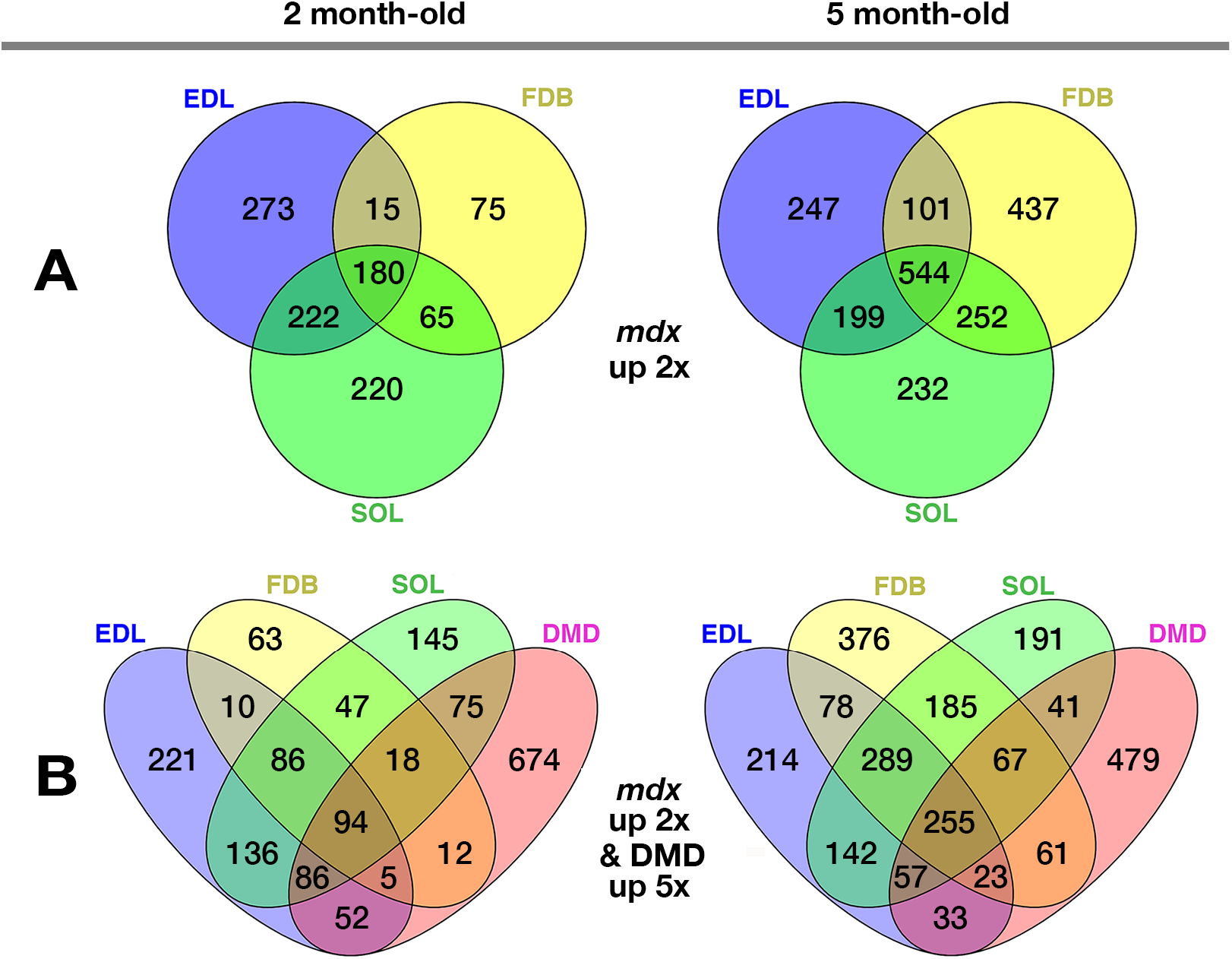
ANOVA comparison of mdx vs. WT and mdx vs. DMD muscles shows a large effect of muscle type and age. ANOVA comparisons of *mdx* vs. WT and DMD vs NORM gene expression were performed as described in Methods 2.4 and 2.5. Numbers of differentially expressed (DE) genes were then compared across conditions in Venn diagrams. A: Comparison of DE genes that were up ≥ 2-fold in all three muscles of 2- or 5 month-old mice; B: Similar to A but including DE genes that were up ≥ 5-fold in DMD vs. NORM patients. The more than 2-fold increase with age in genes common to all samples (180 to 544 in A; 94 to 255 in B) is mainly due to an increase in FDB DE genes. A list of the genes can be found in the Supplementary Material file (SuppTable1).

### 3.2. Developmental and structural as well as inflammatory and immune pathways are enriched in mdx muscles

Pathway enrichment analysis was then performed in Metascape.org. The software integrates the results across conditions by hierarchical clustering. The resulting plot (Fig. 4) highlights the diversity of pathways affected. These include inflammatory and immune pathways but also developmental and structural pathways such as negative regulation of cell proliferation, extracellular matrix organization, actin filament-based processes, MAPK cascade and others. Furthermore, the plot highlights age- and muscle-related differences. All 2 month old groups show higher activation of inflammatory and immunological than of other pathways but this is, again, much more noticeable for the FDB. At 5 months, in contrast, the three muscles show quite similar differences between *mdx* and WT genotypes.

**Figure 4:**
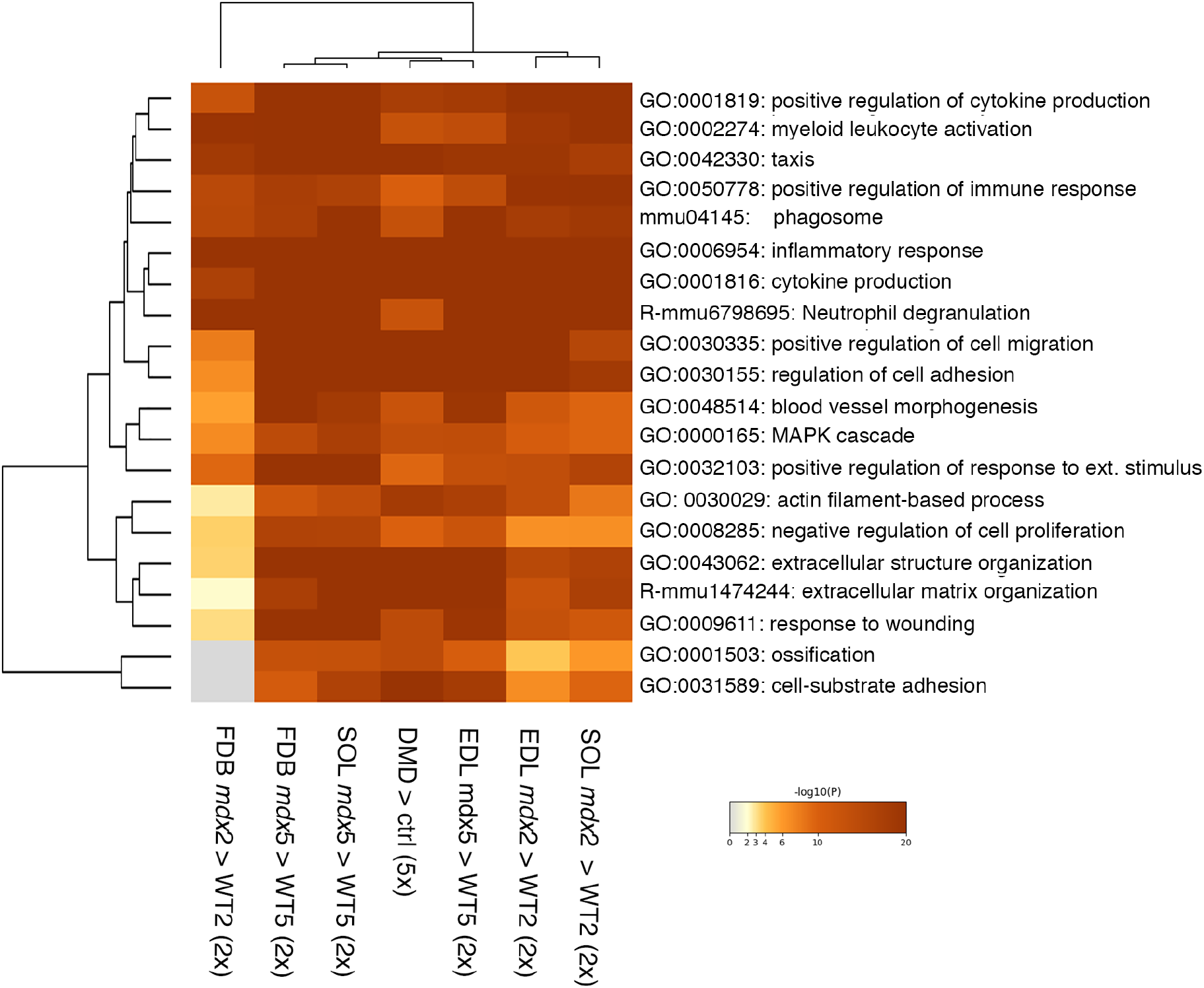
Common metabolic, structural and developmental pathways are affected in both mdx and DMD muscles. Genes differentially expressed in *mdx* compared to WT muscles of both 2- and 5- month-old mice were compared to those in DMD muscles from subjects aged 11 months to 8 years (see Methods 2.5). Pathway enrichment analyzed in Metascape is expressed as −log (P). Pathways related to inflammation and immune response are activated in each mouse and human sample group. Other pathways are affected as well in human and 5 month-old mouse muscles but to a lesser degree in 2 month-old SOL and EDL and practically not at all in 2 month-old FDB.

### 3.3. The mdx mouse is a model of early DMD changes

We next asked whether the relatively mild pathology of the *mdx* mouse may simulate the early stages of DMD. To do so we compared our mouse ANOVA results with two earlier microarray analyses of DMD: a study of 5- to 7-year-old DMD boys [38] and a study of younger than 2-year-old, pre-symptomatic DMD boys [39]. Together these two studies led to the definition of an “early molecular DMD signature” [39]. We also included our ANOVA analysis of the RNA-seq analysis [1] of DMD patients of 11 months to 8 years of age, with mild to severe histopathology. The complete results are presented in Table S2. Genes are grouped by categories as previously defined [38]. We had to remove 13 genes (listed in Table S2) whose names differed between the Haslett and Pescatori publications [38–39] and could not be reconciled. Furthermore, three genes (*CFHR1*, *TRO* and *MEG3*) could not be found under this or any other associated name in the Khairallah study [1]. The presentation as a bubble plot (Fig. 5) shows the agreement not only between the three human studies (indicated as H for Haslett, P for Pescatori and K for Khairallah) but also between these and our mouse work. Each bubble represents the DMD vs. NORM or *mdx* vs. WT comparison for one gene, muscle & age. Color reflects the fold change; diameter reflects the significance, as indicated on the figure. For the more than 80 “disease signature genes” in this figure, expression is either increased or decreased consistently across human and mouse datasets. The mouse data show an age effect: except for 10 genes, the *mdx* vs. WT fold change is smaller or less significant at 2 than 5 months and this is true for each of the 3 muscles. We conclude that the *mdx* mouse transcriptome shares with DMD an early disease signature better manifested in the 5- than 2- month-old mice.

**Figure 5:**
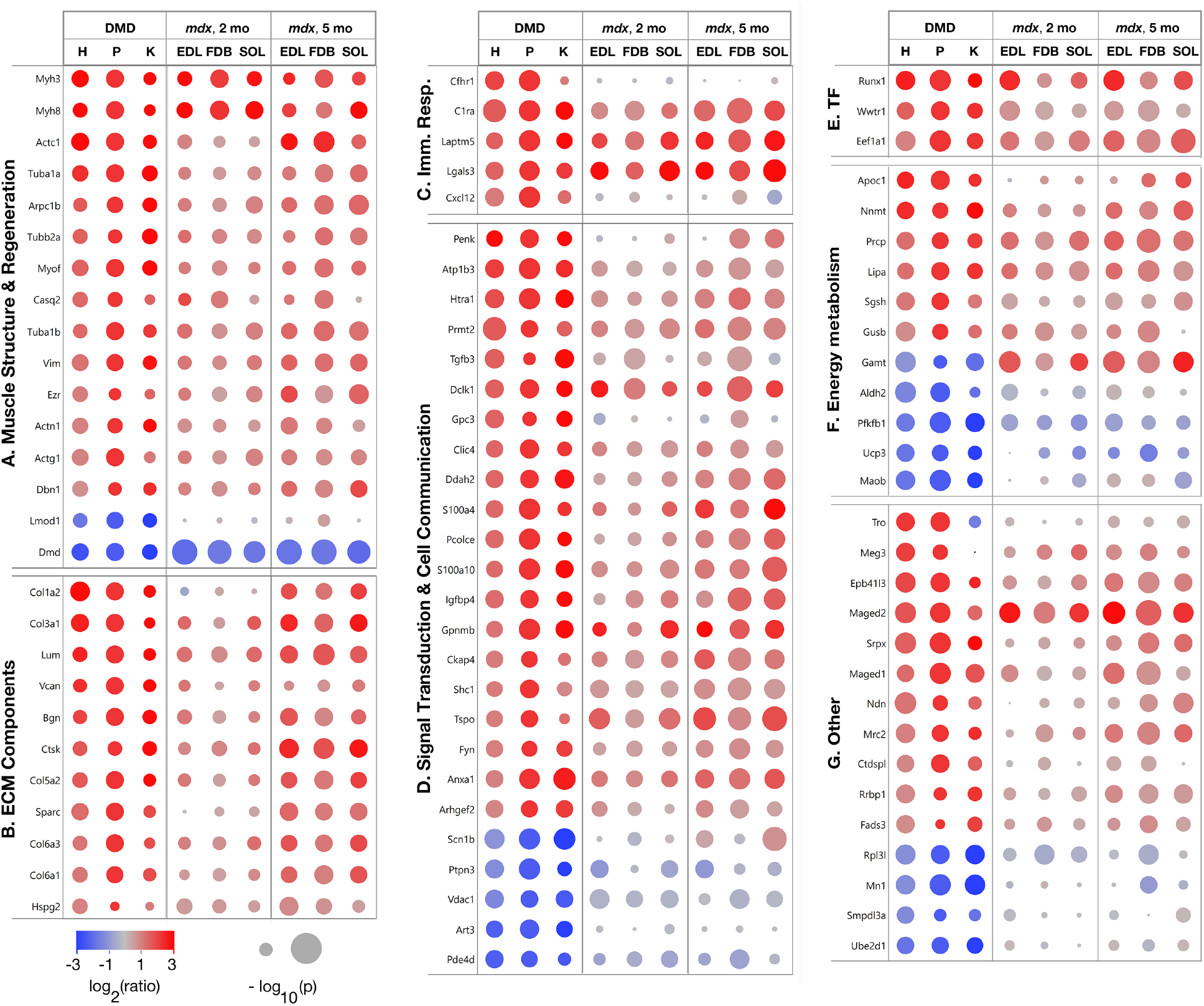
A large set of genes that forms a “molecular signature” of DMD, is similarly affected in the mdx mouse. This figure presents a bubble plot for visualization of the results presented in SuppTable 3. Genes are organized in 7 categories as they were in the original work of Haslett et al. [38] and Pescatori et al. [39]. Human biopsies in both were collected from the *quadriceps* muscle.Their data are in column H and P respectively. Column K contains data from Khairallah et al. [1] recalculated by us from their original log_2_ (RPKM) data. Bubble colors reflect the fold difference DMD vs. NORM or *mdx* vs. WT for the indicated gene and mouse age groups. Bubble diameters are proportional to the significance of the result, representing −log (p).

### 3.4. Analysis of myosin heavy chain isoforms does not support the notion of major differences in muscle fiber type between mdx and WT muscles

Since gene expression in the three muscles investigated remains distinct regardless of age and genotype (Figs. 1 and 2), they probably do not undergo major changes in fiber type. However, DMD may affect fast fibers preferentially [44–45] and the *mdx* SOL muscle is protected, compared to the EDL, from the force loss caused by eccentric contractions [36]. Among the proteins linked to this protection are the myosin heavy chains MyHC2, MyHC4 and MyHC7 which are characteristic, respectively, of fiber types 2A (fast intermediate), 2A (fast), and 1 (slow), respectively. We calculated the differences in their gene (*MyH*) isoform expression in the different mouse groups and the proportion of each fiber type in the two genotypes and ages (Fig. 6 and Table S3). The only highly significant changes are those of *MyH3* and *MyH8*, the already mentioned regeneration-linked isoforms. ANOVA analysis comparing SOL to EDL (Table S4) found differences in the Serca1 Ca^++^-ATPase gene *Atp2a1* and in *MyH2*, *MyH4*, and *MyH7* but not in the cytoplasmic β- and γ-actins and utrophin, also implicated in eccentric contraction protection [36]. Both transcriptional and non-transcriptional pathways must be involved in the SOL protection.

**Figure 6:**
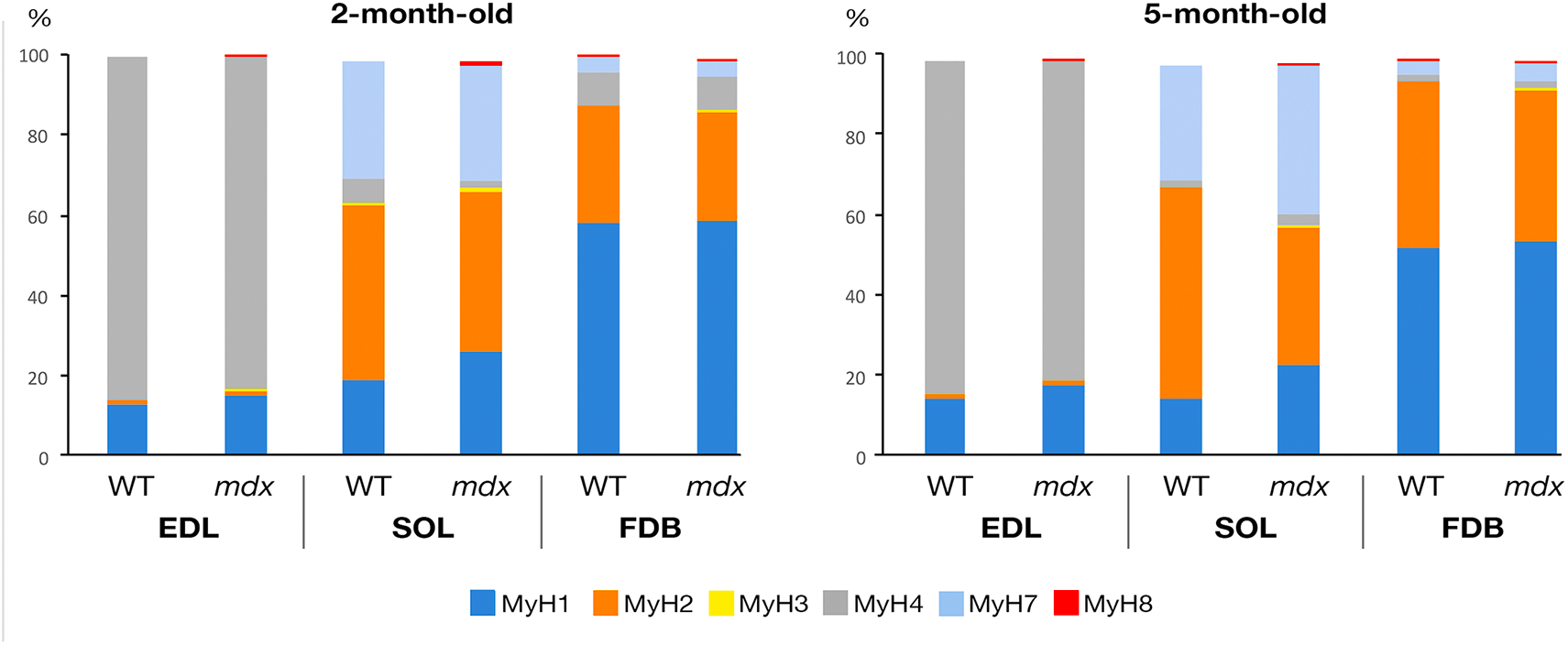
Changes in the proportion of myosin heavy chain (MyHC) isoform gene expression between WT and mdx samples show modest disease-related changes except for minor forms characteristic of muscle regeneration. From the ANOVA results, the proportion of each *MyH* gene was calculated in % of the total and the ratio and p-values were calculated (Table S3). Only those isoforms that contribute at least 1% are included in the bar graphs. The major isoforms are *MyH1* (characteristic of fast 2X fibers [37]), *MyH2* (fast 2A), *MyH*4 (fast 2B) and *MyH7* (slow I). The minor isoforms *MyH3* (embryonic) and *MyH8* (perinatal) are barely noticeable in the graph but their *mdx* vs. WT fold increase is the highest of all isoforms; 24.9 for *MyH8* in the SOL, 18.6 for *MyH3* in the EDL. The values are lower for the FDB (5.1 for *MyH8*). These differences are also most significant. They signal contributions of muscle regeneration.

### 3.5. Human muscle fibers do show a grid-like organization of microtubules

Experimental eccentric muscle contractions also cause a rapid loss of transverse microtubules in mouse muscles [46], a striking correlation between microtubule organization and muscle function. However, we had so far not been able to visualize human muscle microtubules for the lack of properly fixed and maintained human fibers. Here we show immunofluorescent staining of human muscle fibers for α-tubulin (Fig. 7A, in magenta). This confocal image, captured at the surface of the fiber, shows a grid-like network of longitudinal and transverse microtubules characteristic of fast fibers [47]. Double staining with anti-GLUT4 (Fig. 7A, in green) marks the trans-Golgi network [41]. White arrowheads indicate microtubule nucleation centers [32]. Microtubule directionality (Fig. 7B) was analyzed on 10 images from 4 different human fiber donors by the TeDT software ([43] and Methods). The presence of peaks at 0 and 90 degrees confirms the presence of an orthogonal grid of microtubules. We thus conclude that human muscle microtubules have a similar organization to that of mouse and rat muscle and may play a role in the damaging effects of eccentric contractions on DMD patients.

**Figure 7:**
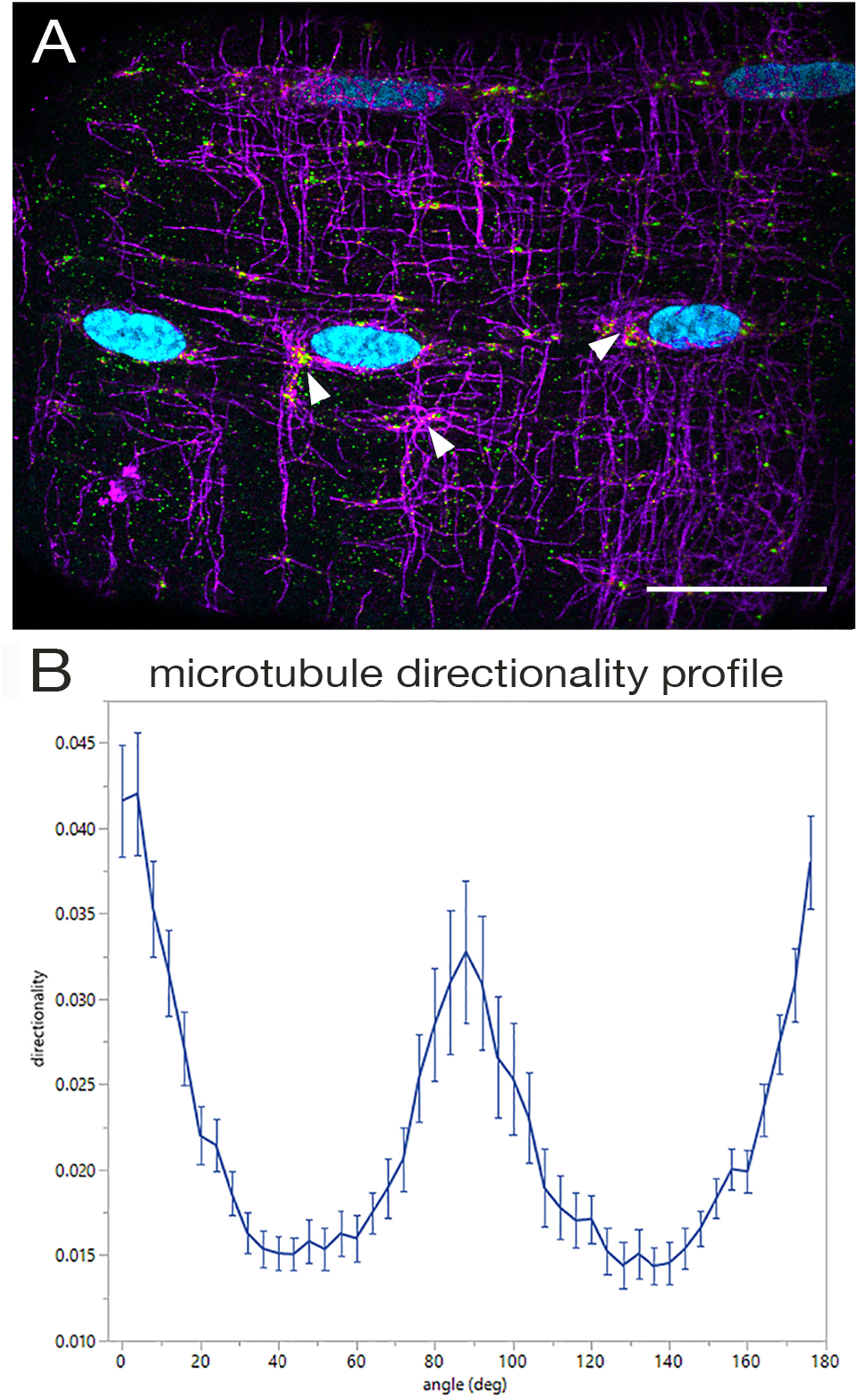
Human muscle microtubules form a grid similar to that of rodent muscle microtubules. Rodent muscle microtubules form a grid that has been linked to the health status of the muscle [1] but, to the best of our knowledge, human muscle microtubules have not been previously visualized. ***A***: Single teased fibers prepared from muscle biopsies from healthy donors were double-stained (see Methods 2. 8.) with an anti-human alpha-tubulin antibody to label microtubules (shown in magenta) and with an anti-human GLUT4 antibody to label trans-Golgi elements (shown in green) that are involved in microtubule nucleation in muscle. Nuclei were counterstained with Hoechst 33342 (shown in cyan) and images were collected with an SP5 Leica confocal microscope. Arrowheads point to some of the microtubule nucleation centers. Bar: 25 micrometers. ***B***: The data for the directionality plot was calculated with the TeDT software ([43] and Methods 2.9.) and plotted in jmp14. The plot represents the average directionality of microtubules in 10 such fibers, plus or minus the standard error. The presence of microtubules at 0/ 180 degrees (i.e. aligned with the longitudinal axes of the fibers, and of other microtubules at 90 degrees (i.e. transverse) confirms the presence of an orthogonal grid in human fibers.

## 4. DISCUSSION

The *mdx* mouse has been criticized for its mild pathology [23] but it remains in use and there is practically no new attempt at curing DMD that is not first tested on the *mdx* mouse [30]. Here we show that the *mdx* muscle transcriptome, analyzed in three different hindlimb muscles, reflects the transcriptome of presymptomatic human DMD muscles, revealing a “disease signature” [38–39], consistent with the mild *mdx* pathology. We found the overall consistency between three sets of earlier human DMD data (micro-array data from 2002 and RNA-seq data from 2012) overall comforting. Some genes that show strong differences between *mdx* and DMD results turned out to be predominantly expressed in non-skeletal muscle tissues. Among these are *Lmod1* (mostly expressed in smooth muscle), *Cxcl12* (mostly in immune cells), *Penk* (much higher in brain than muscle), *Smpdl3a* (mostly in blood serum and macrophages) and *Ube2d1* (mostly in bone marrow and heart). The proportion of these tissues or cells may increase in DMD muscle biopsies as the pathology progresses. However we cannot rule out real human vs. mouse differences which we did not investigate.

The present results validate using the FDB muscle for DMD studies, since similar pathways are enriched in the FDB as in the larger hindlimb muscles, although the differences between *mdx* and WT muscles build up more slowly in the FDB compared to the EDL and SOL muscles (Fig. 5 and Table S3). A few genes nevertheless are more highly expressed at 2 than at 5 months, for example *MyH3* and *MyH8*, two myosin heavy chain isoforms whose increased expression indicates muscle regeneration. They are among the five genes with most increased expression in the DMD vs. NORM RNA-seq, but in the *mdx* mouse regeneration does not continue to increase after an initial burst at 4-6 weeks [48]. A proteomic study carried out on 3 month old mice found a lower proportion of regenerating muscle fibers in the FDB than in the EDL *mdx* muscles at that age [49]. The SOL has been reported to be most similar to human muscles [50].

RNA-seq data are very useful in the parsing out of a tissue’s response to pathological conditions but many genes are not transcriptionally regulated or may undergo more than one mode of regulation. The cytoplasmic actins that are implicated in the muscle response to eccentric contraction [36] showed a minor transcriptional response, in agreement with post-transcriptional and post-translational regulatory pathways [51], also regulating tubulins [52] and utrophin [53].

Microtubule organization is a sensitive marker of muscle health, likely because it is regulated by patterned activity of the muscle [54]. It could in principle serve as a biomarker but this has not been possible. First, attempts to find a correlation between microtubule directionality and physiological parameters in *mdx* mouse lines rescued by partial DMD constructs did not give straight results [17]. There is steady progress, however, as a link has been made between the presence of transverse microtubules and the effect of eccentric contractions [46]. Secondly, while it is possible to find normal human fibers fixed and suitably treated [53] to permit immunofluorescent staining of microtubules, it has been impossible, so far, to find equivalent DMD samples. DMD muscle biopsies, when collected, are immediately frozen. Regrettably, microtubules do not survive the freeze-thaw. We therefore do not know what DMD muscle microtubules look like.

The newer mouse lines developed to improve on *mdx* are more severely affected but do not cover each aspect and stage of DMD either. For example, the D2-*mdx* mouse, which has the DBA/2 instead of the C57BL/10 genetic background [29, 55], may be useful for investigating fibrosis since D2-*mdx* shows fibrosis in the diaphragm and in hindlimb muscles while *mdx* shows fibrosis in the diaphragm only [14]. Several new animal lines seem to be better models of the cardiac than of the skeletal aspects of DMD: the Cmah−/−;-*mdx* mouse, which carries a human-like mutation in a gene involved in sialylation [28, 56]; the *mdx*/mTR mouse [27, 57] which has shorter telomeres than the *mdx* mouse, as in the human disease; and a new *mdx* rat model [58]. Different mouse lines have been compared for studies of bone defects [59] which the *mdx* mouse does not mimic well [60] although it has been used successfully to study pharmacological effects on bone [61]. Finally, the GRMD, the dog line that best models DMD still diverges from the human disease and does not always show a severe pathology and shortened life. It remains a challenge to find the best animal model for DMD [10,62]. In the meantime, the *mdx* mouse remains essential. Our RNA-seq analysis is available to the public as is the RNA-seq analysis of DMD [1]. Together they should help pinpoint transcriptional and post-transcriptional contributions to the early development of DMD and allow a better evaluation of the contribution of animal age in such studies. Finally, we add a missing link to the muscle microtubule story by showing that human muscle microtubules, like those of rodents, are organized in a grid pattern that appears essential for muscle health.

## ACKNOWLEDGEMENTS

This study was supported by the Intramural Program of the National Institute of Arthritis and Musculoskeletal and Skin Diseases and utilized the computational resources of the NIH HPC Biowulf cluster (http://hpc.nih.gov). We are grateful to Drs. Hong-Wei Sun (Biodata Mining & Discovery Section, NIAMS), Olivier Duverger (NIDCR) and Davide Randazzo (NIAMS) for helpful discussions, to Dr. Thorkil Ploug and the FINE team (University of Copenhagen, Denmark) for the gift of human muscle fibers, and to Drs. Jim Ervasti (University of Minneapolis, MN) and Alessandra Pasut (Katholieke Universiteit Leuven, Belgium) for critical reading of the manuscript.

## Footnotes

1 ***Abbreviations:*** DMD: Duchenne muscular dystrophy; DE: differentially expressed; EDL: *extensor digitorum longus;* FDB: *flexor digitorum brevis;* GLUT4: Glucose transporter type 4; GRMD: Golden Retriever dog model; *MyH*: myosin heavy chain (gene); MyHC: myosin heavy chain (protein); NORM: normal; PCA: primary component analysis; ROS: reactive oxygen species; RPKM: reads per kilobase of transcript, per million mapped reads; SOL: *soleus;* WT: wild type.

