## Supplementary material for "Transcriptomic analysis of mdx mouse muscles reveals a signature of early human Duchenne muscular dystrophy": Tables S1-S4 Legends

***SUPPLEMENTARY MATERIAL LEGENDS***

***Table S1: Differentially expressed genes plotted as Venn diagram in Figure 3***

ANOVA analysis was carried out to compare *mdx* vs WT and DMD vs NORM groups. Differentially expressed genes were selected using a cutoff of 2-fold increased expression for mouse groups and 5-fold increased expression for human groups.

***Table S2: The change in expression of "disease signature" genes, initially identified in samples from presymptomatic DMD boys, is also observed in mdx mouse muscles.***

The Table lists 86 genes in 7 different categories. There were initially 105 genes presented by Haslett et al. [38] as increasingly expressed in muscle microarrays with samples from 5-7 year old DMD boys. Pescatori et al. [39] found a significant difference and increase in expression for 86 of these genes in samples from presymptomatic 1-2 year old DMD boys and coined the term "disease signature" genes. In this table we compare these original results to our mouse data. We also add the more recent RNA-seq results from human samples of different ages and DMD stages [1] after our own re-analysis of these results. For each sample, the results presented are p-value and DMD vs. normal or *mdx* vs. WT fold expression change. Non-significant changes (p>0.05) are shown in gray, significant changes are shown in red (up) or blue (down). See also Figure 6 and main text.

***Table S3: Changes in myosin heavy chains (MyH) between WT and mdx muscles are generally modest, except for isoforms that reflect muscle regeneration.***

Table A provides the fold change and p-values for changes in MyH expression between WT and *mdx* muscles. For the main MyH isoforms (MyH1, MyH2, Myh4 and MyH7) the fold changes are generally modest. In contrast, for the minor MyH isoforms MyH3 and MyH8 which are expressed in embryonic and perinatal muscle fibers and thus are increased during regeneration, the fold changes are larger and highly significant emphasizing muscle regeneration.

Table B gives the proportion of MyH (in %) calculated from the RPKM data for each mouse muscle, genotype and age. Mouse muscles contain several fiber types; no muscle is 100% fast or 100% slow.

***Table S4***: ANOVA was carried out to compare gene expression in SOL vs. EDL muscles. This table shows the results for genes implicated in the protection of the slow SOL compared to the fast EDL muscle from force loss following eccentric contractions [36. Some of the genes implicated in this force loss (Atp2a1 and the three MyH e.g.) show large and highly significant expression differences between the 2 muscles. In contrast, utrophin shows no significant difference, in agreement with data indicating that it is post-transcriptionally regulated [53]. The cytoplasmic actins as well show only very small differences between the muscles.
